# ADRA2B and HTR1A: An updated study of the biogenic amine receptors reveals novel conserved motifs which play key role in Mental Disorders

**DOI:** 10.1101/2022.09.16.508280

**Authors:** Louis Papageorgiou, Evangelia Christou, Effrosyni Louka, Eleni Papakonstantinou, Io Diakou, Katerina Pierouli, Konstantina Dragoumani, Flora Bacopoulou, George P Chrousos, Elias Eliopoulos, Dimitrios Vlachakis

**Author notes:** Correspondence to*: Dr Dimitrios Vlachakis, Laboratory of Genetics, Department of Biotechnology, School of Applied Biology and Biotechnology, Agricultural University of Athens, 75 Iera Odos, 11855 Athens, Greece.

## Abstract

Mental disorders are strongly connected with several psychiatric conditions including depression, bipolar disorder, schizophrenia, eating disorder and suicides. There are many biological conditions and pathways that define these complicated illnesses. For example, eating disorders are complex mental health conditions that require the intervention of geneticists, psychiatrists and medical experts in order to alleviate their symptoms. A patient with suicidal ideation should first be identified and consequently monitored by a similar team of specialists. Both genetics and epigenetics can shed light on eating disorders and suicides as they are found in the main core of such investigations. In the present study, an analysis has been performed on two specific members of the GPCR family towards drawing conclusions regarding their functionality and implementation in mental disorders. Specifically, evolutionary and structural studies on the adrenoceptor alpha 2b (ADRA2B) and the 5-hydroxytryptamine receptor 1A (HTR1A) have been carried out. Both receptors are classified in the biogenic amine receptors sub-cluster of the GPCRs and have been connected in many studies with mental diseases and malnutrition conditions. The major goal of this study is the investigation of conserved motifs among biogenic amine receptors that play an important role in this family signaling pathway, through an updated evolutionary analysis and the correlation of this information with the structural features of the HTR1A and ADRA2B. Furthermore, structural comparison of ADRA2B, HTR1A, and other members of GPCRs related with mental disorders is performed.

## Introduction

Eating disorders are characterized as mental disorders defined by abnormal eating habits that affect negatively patient’s physical or mental health. The most famous of them are anorexia nervosa and bulimia nervosa. In anorexia nervosa people avoid eating due to the fear of gaining weight even though they have below average body weight. In bulimia nervosa patients eat a lot and then they try to rid of the food consumed. Mental disorders like anxiety, depression and substance abuse are common among people dealing with eating disorders. Worldwide, anorexia nervosa affects about 0.4% and bulimia nervosa affects about 1.3% of the population of young women per year (1). Eating disorders typically begin in adolescence or early adulthood. The causes of eating disorders are not clear. Both biological and epigenetic factors appear to be a primary cause. The idealization of thinness is believed to contribute to some eating disorders. Treatment involves psychological support, a proper diet and a normal amount of exercise and can be effective for many eating disorders (1). Hospitalization may be needed in more serious cases. Unfortunately, both anorexia nervosa and bulimia nervosa have a high risk of death.

Emotional intelligence is a term referring to a group of non-cognitive abilities accounting for how people accommodate and adapt to intra - and interpersonal conditions by identifying emotions, incorporating emotions in thought processes, understanding emotional complexity, and manipulating emotions in self and others (2). Low emotional intelligence (EI) and emotional dysfunction may have an important role in the development and maintenance of maladaptive eating behavior than is currently emphasized within the biologically based to the forming idea that is often focused within the literature (3). Emotional difficulties have been observed in individuals with disordered eating patterns in a big number of studies, including poor consciousness, difficulties with emotional language and confusion of emotional states. These difficulties with emotional functioning are a significant factor related to the core etiology of eating disorders (4, 5). However, limited knowledge exists to how this impact on professional ability to engage patients within treatment as a result of such dysfunction (3). As Faye, Hazlett *et al*. found in their study, aspects of EI such as emotional regulation and lack of an emotional language are considered to be at the core of the onset and maintenance of these disorders.

Many patients before the onset of the disorder report more social difficulties, fewer childhood friends, and engage in more solitary activities than healthy controls (6). During eating disorders, a variety of difficulties are seen, such as poorer social skills and social problem - solving abilities, high social anxiety, reduced social networks, and loss of interest in social activities. So, socio-cognitive mechanisms have an important role in eating disorders. Furthermore, social appearance anxiety which is the fear when an individual perceives that his or her body is not meeting the ideal body type accepted by a particular culture, leads individuals to experience body shame and dissatisfaction, social appearance anxiety, and other negative effect, which may in turn lead to disordered eating patterns (7). Several studies suggest that lower emotional intelligence increases the likeliness of eating disorders (2, 8). According to the individuals interviewed in a study (3), patients use maladaptive eating patterns as a way to deal with difficult life situations and experiences.

On the other hand, suicidal behavior is considered as a paradoxical human behavior that arises as a product of complex mental and emotional processes. Suicidal behavior can be explained in a percentage of 43% by genetics and the remaining 57% by environmental factors. Suicide is a complex, multifactorial phenomenon, with specific biological, psychological and environmental risk factors, including: Psychiatric disorders (mainly depression), previous suicide attempt, psychosocial factors (stressful life events), personality traits (impulsivity and aggression), history of traumatic experiences and / or abuse, family history of psychiatric disorders and suicidal behavior, substance and alcohol abuse and general medical conditions (9). It is known that mental disorders account for the vast majority of suicides and suicide attempts. The numbers are at least 10 times higher than the general population. The reported rate of completed suicides in this context varies between 60% and 98% of all suicides. Many of the other incidents are related to financial, relationship and related issues. However, other causes are discrimination, violence and war as well as eating disorders (10, 11). Suicide is a global phenomenon. According to the World Health Organization, in 2015, approximately 800,000 suicides were recorded worldwide, and worldwide 78% of all suicides occur in low- and middle-income countries. Overall, suicides account for 1.4% of premature deaths worldwide, and it is noteworthy that the number of suicides in the world has risen sharply among young people. Indeed, suicide is the second leading cause of death for adolescents aged 15 to 19 years (10, 12). Regarding the rates between the two sexes, there is what is known as the gender paradox in suicide: women make more suicide attempts than men, as more women than men are diagnosed with depression, but success rates are higher in men (4: 1 in most developed countries). This may be due, at least in part, to differences in the gender roles that correspond to men and women which can lead to stress. A correlation has yet been found between the violence experienced by a woman by a familiar person (intimate partner violence, IPV) and suicidal ideation and efforts (13, 14).

The past decade has been significant progress to understand the genetic and epigenetic effects of eating disorders and suicide. Eating disorders often occur in families with a history background and studies on twins have revealed that genetic factors account for about 40Po to 60a of the causes of anorexia nervosa, bulimia nervosa, and other eating disorders. Similarly, suicidal behavior accumulates in families. Twins and adoption studies suggest that heredity is 30 to 50%, although a more accurate estimate could be 17 to 36% if the heredity of comorbid psychiatric disorders is checked. Molecular genetic studies have also been performed to detect alterations in DNA sequence or in gene expression that may be involved in the pathogenesis of eating disorders and their symptoms on the one hand and suicidal behavior on the other. It has been suggested that there is a link between eating disorders, polymorphisms of the brain-derived neurotrophic factor (BDNF) and serotonin genes. In particular, the Val66Met polymorphism of the BDNF gene has been associated with anorexia nervosa and bulimia nervosa has anorexia as well as schizophrenia and depression (15, 16). In addition, significant correlations have been found between the allele A (and the homozygous A / A genotype) and the 102T/C polymorphism in 5-HT2A serotonine receptor with NA/N B and suicidal ideation in patients with major depression, respectively (17). Finally, a recent meta-analysis maintains that there is a link between the 5- HTTLPR gene mutations and anorexia nervosa, as well as violent suicidal behavior, but not with bulimia nervosa (18). Generally, there are many studies which suggest that there is a link between eating disorders and the pathway of the gene expression of serotonin, catecholamine, neuropeptide and feeding regulation (19, 20). Suicidal behavior and most of the eating disorders are highly connected with mental issues such as depression and hormone expression like serotonin and dopamine. This may be a reason why eating disorders and many mental disorders are affected by the biological pathways of hormones expression and neurotransmitters.

G protein-coupled receptors (GPCRs) regulate physiological functions to maintain homeostasis by mediating most of our physiological responses to hormones (mainly catecholamines), neurotransmitters and environmental stimulants and so have great potential as therapeutic targets for a broad spectrum of diseases. The family of G- protein-coupled receptors (GPCRs) is categorized in five main families, named glutamate, rhodopsin, adhesion, frizzled/taste2, and secretin, forming the GRAFS classification system (21). All GPCRs are characterized by the presence of seven membrane-spanning a-helical segments which are separated by alternating intracellular and extracellular loop region (Figure1) (22–24). The extracellular loops and N termini of GPCRs, together with the extracellular halves of the transmembrane helices, are believed to define the ligand-binding site of each receptor. Therefore, the extracellular loops may play an important role in the overall pharmacology of any particular receptor (25).

**Figure 1.**
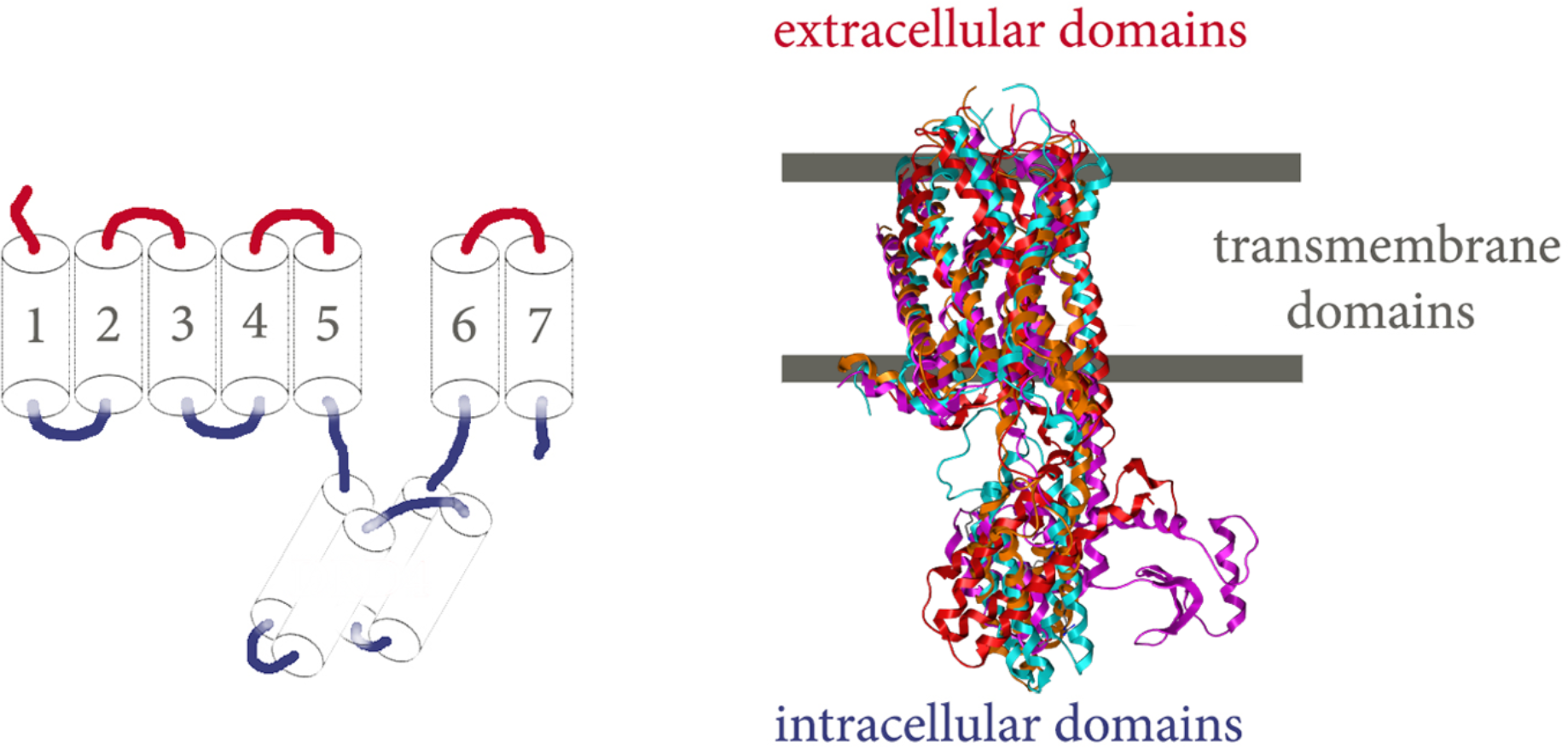
Structural architecture of biogenic amine receptors family receptors based on the available 3D structures from the Protein Data Bank. A. Cartoon-like description of the main structural architecture of biogenic amine receptors. The seven transmembrane helices are numbered, the extracellular domains are colored red and the intracellular domains blue. B. Comparison of DRD2 (purple), HTR1B (blue), DRD4 (red) and ADRA2A (orange) 3D structures from the Protein Data Bank (PDB: 6CM4, 4IAR, 5WIV and 6KUY). The transmembrane area is colored grey, the extracellular domains are colored red and the intracellular domains blue.

The main difference between the members of GPCRs is the domain which is out of the membrane and in case of both ADRA2B and HTR1A is an intracellular domain (Figure 1 and 2). During activation, the GPCR opens a cavity on its cytoplasmic side, between the fifth and the sixth helix and the remaining of the receptor, to interact with downstream effectors such as G-proteins andarrestins (26–28). Based on several phylogenetic analyses, G protein-coupled receptors have been categorized into several families on the basis of their sequence and structural similarity. The A-family or rhodopsin-like is the largest one including over 700 members from human genome alone, where both adrenergic and 5-hydroxytryptamine receptors are classified. This family possesses a number o1 highly conserved characteristic motifs which may play important structural and functional roles in signaling mechanisms which are shared by all members of their family (29–34).

**Figure 2.**
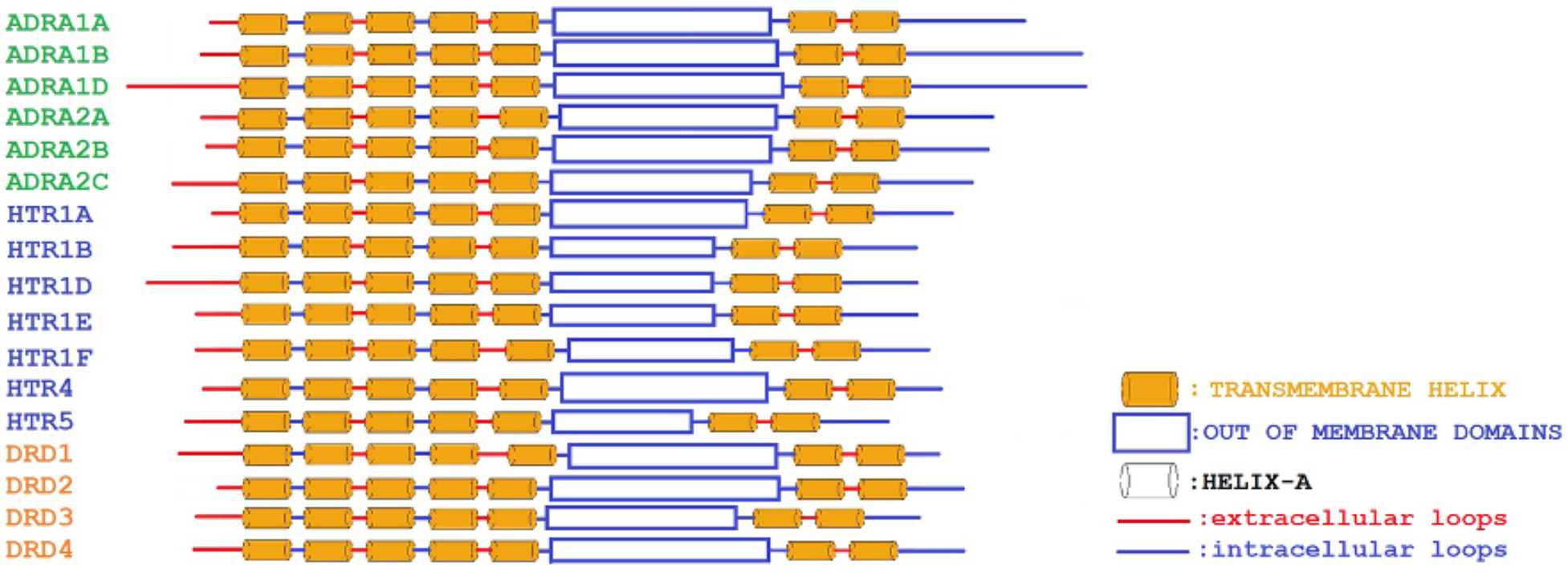
Cartoon like architecture of selected types of biogenic amine receptors based on the Interpro. There are presented three different types and sub-types of biogenic amine receptors ADRA (colored green), HTR (colored blue) and DRD (colored orange). We compared the extracellular loops (lines colored red), the intracellular loops (lines colored blue), the transmembrane helices (barrels colored orange) and the intracellular domains (rectangles colored blue). From this comparison we found out that there are many differences mainly at the extra- and intra-cellular loops and the intracellular domains of all the members.

One important group of neurotransmitters that belong to the GPCR family is biogenic amine receptors. These molecules interact with endogenous bioamine ligands, such as dopamine, serotonin, and epinephrine, are targeted by a wide array of pharmaceuticals. They exert significant influence on both behavior and physiology. The biogenic amine receptor family, which includes dopamine (DRD), histamine (HRH), trace (TAAR), adrenergic (ADR), muscarinic cholinergic (mAChR), and most serotonin (5HTR) receptors, primarily mediates biogenic amine activity (35). Like all GPCRs, biogenic amine receptors have a characteristic structure of seven transmembrane domains separated by three extracellular and three intracellular loops (Figure 1 and 2). They propagate intracellular signaling through a G protein-mediated pathway. Furthermore, these receptors are well-known protein targets against myriad diseases such as migraines, hypertension, schizophrenia, asthma, allergies, and stomach ulcers.

Adrenergic receptors (or adrenoceptors) are rhodopsin-like G protein-coupled receptors and also belong to the biogenic amine receptors family. They are classified in two groups, alpha (ADRA) and beta (ADRB), and there are several subtypes in each group. Within the alpha-type, the a2-adrenoreceptors (ADRA2) are composed of three subtypes: ADRA2A, ADRA2B and ADRA2C (36). They respond to catecholamines’ line, especially norepinephrine (noradrenaline) and epinephrine (adrenaline) and couple to the Gi family of G proteins. Many cells possess these receptors, and the binding of a catecholamine to the receptor will generally stimulate the sympathetic nervous system, effect blood pressure, myocardial contractile rate and force, airway reactivity, and a variety of metabolic (for example insulin secretion and lipolysis) and central nervous system functions. The clinical uses of adrenergic compounds are varied.

Adrenoceptor alpha 2b is an intron less gene that encodes a seven-pass transmembrane protein. This protein regulates neurotransmitter release from sympathetic nerves and from adrenergic neurons in the central nervous system. Alpha-2 adrenergic receptors mediate the catecholamine-induced inhibition of adenylatecyclase through the action of G proteins (36).The whole 3D structure of ADRA2B has not been well predicted yet apart from some sections of it. Especially the intracellular domain of ADRA2B is still remaining unsolved. Recent studies have predicted the transmembranic domain of ADRA2B and its active-state by dexmedetomidine (37), but the other parts are still not being defined. This α2BAR—Go complex provides the first active-state structure of adrenoceptors coupled to the Gi/o family of G proteins and explains the molecular origins of subtype selectivity between a- and β- adrenergic agonists. From molecular dynamics simulations have been found that the binding status ofdexmedelomidine in the complex is stable for 2 ps. Dexmedetomidine occupies a position in the binding pocket in the transmembranic domains and interacts with the adrenoceptor alpha -2b primarily through aromatic and van der Waals interactions. The transmembranic domains in adrenergic receptors have some conserved domains that help with the agonist-binding affinity for catecholamines and the activation of the receptor.

Many studies support that a functional deletion variant of the ADRA2B gene encoding the α2B-adrenergic receptor has shown to increase emotional memory and neural activity in the amygdala (38). In particular, this deletion variant is characterized by a loss of three glutamic acids residues (301–303) in the third intracellular loop which encodes the alpha-2b subunit of the noradrenaline receptor (39). As a result, this variant increases noradrenaline availability and as it has been shown carriers have enhanced emotional memory (38). This may justify the reason that ADRA2B relates to mental and eating disorders.

Serotonin receptors are a class of the superfamily GPCRs and ligand-dependent ion channels located in the central and peripheral nervous systems. They mediate both stimulatory and inhibitory neurotransmission. Serotonin receptors are activated by the neurotransmitter serotonin, which acts as their natural ligand. Serotonin receptors regulate the release of many neurotransmitters including glutamate, GABA, dopamine, epinephrine / norepinephrine and acetylcholine, as well as many hormones, such as oxytocin, prolactin, angiopressin, cortisol, corticotropin and substance P, among others. They affect various biological and neurological processes such as aggression, anxiety, appetite, knowledge, learning, memory, mood, nausea, sleep and thermoregulation. They are the target of a variety of pharmaceutical and recreational drugs, including many antidepressants, antipsychotics, anti-obesity, antiemetics, gastro-prokinetic agents, anti-migraine agents, hallucinogens and empathogens. There are seven types of 5 HT receptors, 5 HT1-7, which are further categorized into subtypes. Types I, II, IV, V, VI, and VII belong to the GPCRs except for 5 HT3, which is a ligand gated ion channel (LIC) or ionotropic receptor. Serotonin receptors bind to all three normal signaling pathways via Gai, Gαq / 11 and Gas (G proteins) thus forming various biochemical signaling pathways (40).

The class of 5-HT1 receptors consists of five subtypes of receptors (5 HT1A, 5 HT1B, 5 HT1D, 5 HT1E and 5 HT1F), which are structurally identical in humans at a rate of 40-63%. They bind mainly (but not exclusively) to Gi / G0 proteins. The 5 -HT1A receptor is the most widely distributed of all 5 HT receptors. It is encoded by the HTR1A gene. In addition to serotonin, it acts as a receptor for various drugs and psychoactive substances. Ligand binding causes a change in the configuration that activates signaling through G proteins and regulates the activity of cathodic operators, such as adenylate cyclase. In the brain, 5 HT1A receptors act as auto receptors as well as postsynaptic receptors (41). 5-HT1A is the best studied and is generally accepted to be involved in psychiatric disorders such as anxiety and depression. Both pre- and post-synaptic 5 HT1A receptors play an important role in shaping mood, cognitive function and motivation, functions that are impaired in patients with schizophrenia and are influenced by antipsychotic agents. The discovery of new ligands for the 5 HT1A receptor is the focus of active neurobiological research due to its involvement in psychiatric disorders and even memory loss (42). The 3D structure of 5-HT1A has not been estimated yet. Therefore, there is an urgent need for its structure to be found, as a map that will potentially lead to beneficial knowledge related with a specific structural-based drug design.

## Methods

### Database sequence search

The protein sequences related to the biogenic amine receptors (adrenoceptors alpha, adrenoceptors beta, 5-hydroxytryptamine receptors, dopamine receptors, histamine receptors, trace amine associated receptor and muscarinic acetylcholine receptors) were extracted from the NCBI database. In total, 3259 protein sequences were downloaded and filtered for several species, from which noise data were removed including partial, hypothetical and non-relevant entries. Afterwards, two smaller datasets have consisted of the sub-families of adrenoceptors alpha and 5-hydroxytryptamine receptors.

### Genetic and evolutionary analyses

Multiple sequence alignment (MSA) of biogenic amine receptors including adrenoceptors alpha and 5-hydroxytryptamine receptors were performed using Matlab bioinformatics toolbox (43) utilizing guide trees and the progressive MSA method as previously described in several studies (43–46). Pairwise distances among sequences of each datasets were estimated based on the pairwise alignment with the “Gonnet” method and followed by calculating the differences between each pair of sequences (46).The Neighbor-Joining method was used towards to estimating the guide trees by assuming equal variance and independence of evolutionary distance estimates (45). The phylogenetic analyses for the biogenic amine receptors and also for the adrenoceptors alpha and 5-hydroxytryptamine receptors were performed using the MATLAB Bioinformatics Toolbox utilizing the Unweighted Pair-Group Method (UPGMA) while the matrix of the pairwise distances was calculated using the protein –adapted Jukes-Cantor statistical method (44, 47). The phylogenetic analysis for the biogenic amine receptors was performed using the MATLAB Bioinformatics Toolbox utilizing the Neighbor-Joining method while the matrix of the pairwise distances was calculated using the protein –adapted Jukes-Cantor statistical method (48).

The constructed phylogenetic trees were visualized using MEGA radiation option and the final protein clusters separated in different sub-datasets using a threshold. In total, four major sub-clusters including ADRA1, ADRA2A, ADRA2C, ADRA2B were identified in the phylogenetic analysis of the adrenoceptorsalpha, seven in the phylogenetic analysis of the 5-hydroxytryptamine receptors and nineteen in the phylogenetic analysis of the biogenic amine receptors.

### Specific conserved motifs extraction

The phylogenetic trees that derived from the phylogenetic analyses were separated in sub-trees, in order to extract the most highly related protein sequences of adrenoceptors alpha, 5-hydroxytryptamine receptors and the biogenic amine receptors, respectively for the conserved motifs exploration. Finally, consensus sequences were calculated and visualized for each dataset through JalView platform using the multiple sequences alignment results and parameters include amino-acid conservation (43, 45, 49). The commentary section of JalView, which presents the amino-acid conservation using logos and histograms, was further observed to uncover innovative motifs.

### Molecular modeling of the 5-HTR1A intracellular domain

The amino acid sequence of 5-HTR1A (UniProtKB/Swiss-Prot: P08908.3) with the aid of the blastp algorithm (http://blast.ncbi.nlm.nih.gov/Blast.cgi) was used to identify homologous structures by searching in the Protein Data Bank (PDB). The homology modeling of the 5-HTR1A intracellular domain was carried out using the Molecular Operating Environment version 2013.08 software package developed by Chemical Computing (HPC) cluster (50). The selection of template crystal structure for homology modeling was based on the primary sequence identity and similarity, and the crystal resolution (44, 51). Structural superposition of the candidate template 3D structure has been performed using MOE towards analyzing all the possible conformations cases of the intracellular domain from the GPCRs related the 5-HTR1A. The optimal template has been selected based on the sequence and structural features of available templates and the 5-HTR1A protein sequence.

The crystal structure of the 5-hydroxytryptamine receptor 1B (5-HTR1B) (PDB: 4IAR_A) and the 5-hydroxytryptamine receptor 1A (5-HTR1A) (PDB: 7E2X_R) were used as template structures (37, 52, 53). The MOE homology model method is separated into four main steps. Firstly, a primary fragment geometry specification is performed followed by the insertion and deletion tasks. Consequently, the loop selection and the side-chain packing are performed and the selected final model is refined in the last step. Subsequently, energy minimization was done in MOE initially using Amber99 (44, 54)force-field as implemented into the identical package.

The produced model was initially evaluated within the MOE package by a residue packing quality function, which depends on the amount of buried non-polar side-chain groups and on hydrogen bonding. Moreover, the suite PROCHECK (55) was employed to further evaluate the produced model. Finally, MOE and its implement for protein check module were accustomed to evaluate whether the model of 5-HTR1A domains are similar to known protein structures of this family. The Molecular Dynamics simulations of the 5-HTR1A intracellular domain model was executed in a periodic cell, which was explicitly solvated with simple point charge. The truncated octahedron box was chosen for solvating the models, with a set distance of 7 Å clear of the protein. The molecular dynamic simulations were conducted at 300 K, 1 atm with a set 2 second step size for a total of fifty nanoseconds.

### Structural Comparison

A thorough analysis of GPCRs structural comparisons was performed by superimposing the structures of the Biogenic amine GPCRs amino acids which were closer to the protein-target of the biogenic amine receptors phylogenetic analysis we performed before and the homologous structures the Protein Data Bank (PDB) gave us by searching each receptor. As concerns the 5-hydroxytryptamine receptor 1A the closer amino acids from the phylogenetic tree were HTR1D, HTR5, HTR1B, HTR1E and HTR1F. From the results PDB gave us we selected the crystal structure of the 5-HTR1B (4IAR_A) from Homo sapiens and the 5-hydroxytryptamine receptor 1A (5-HTR1A) (PDB: 7E2X_R).

## Results

### Dataset and Multiple Sequence Alignment

As previous mentioned, the members of the biogenic amine receptors family that we had in our dataset are the sub-families of adrenoceptors, serotonin receptors, dopamine receptors, histamine receptors, trace amine associated receptors and muscarinic acetylcholine receptors. As concerns the serotonin receptors sub-family, 5-hydroxytryptamine receptor 3 is not included in our dataset. The reason is that the receptor 5-HTR3 belongs to the e Cys-loop superfamily of ligand-gated ion channels (LG ICs) and so, differs structurally and functionally from all other 5-HTR (serotonin) receptors which are in the GPCRs family. All the members we have in our dataset are presented in a table (Table 1) where each group is colored with the color we have it in our phylogenetic analysis (Figure 3).

**Figure 3.**
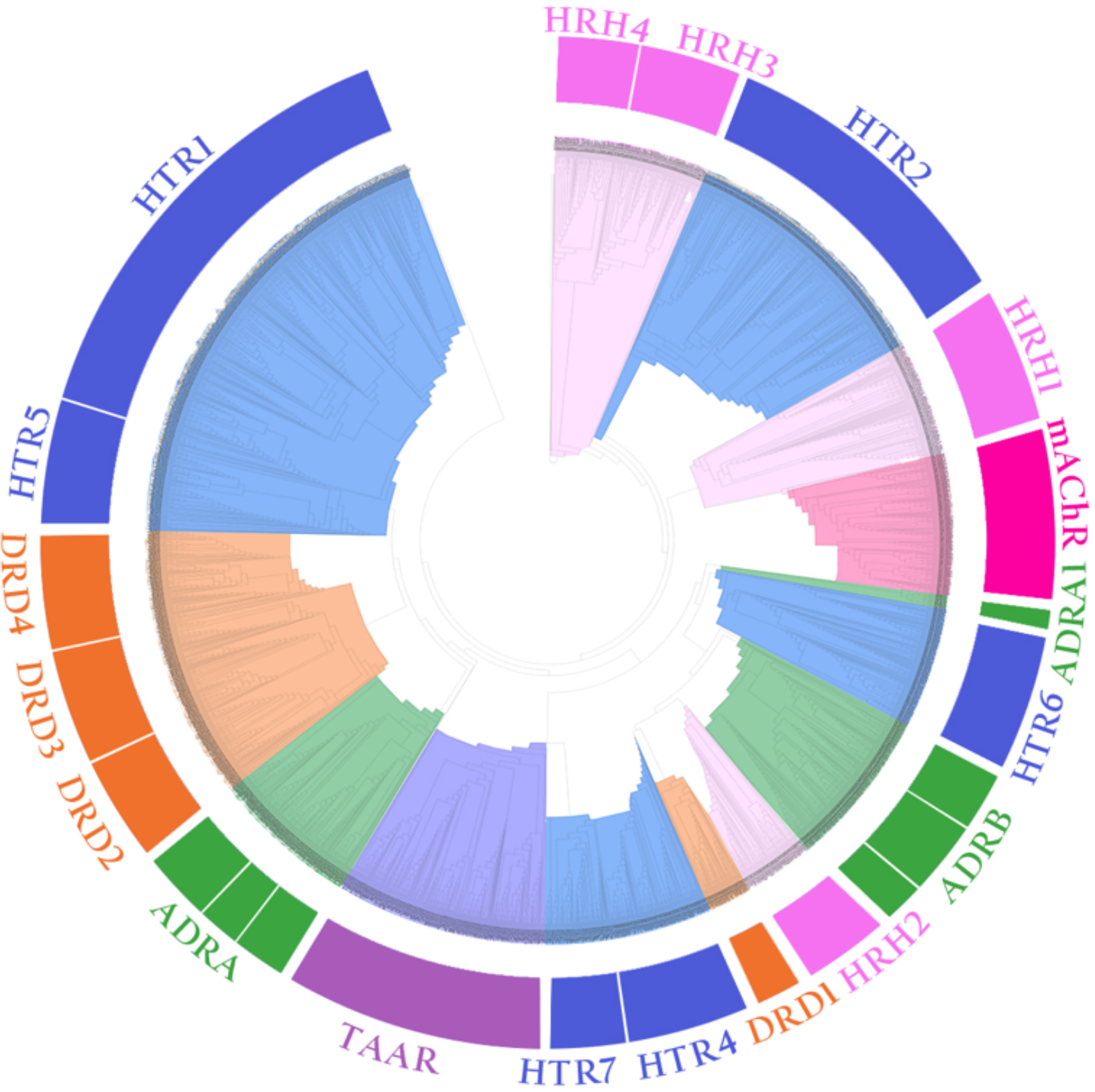
Evolutionary study of the biogenic amine receptors family. The phylogenetic tree was constructed utilizing the Neighbor-Joining method in Matlab Bioinformatics Toolbox and visualized using the TreeExplorer tool of MEGA. In the tree representation there are clearly separated in six branches: HRH (colored pink), HTR (colored blue), mAChR (colored dark pink), ADRA and ADRB (colored green), DRD (colored orange) and TAAR (colored purple).

**Table 1.** Members of the biogenic amine receptors family.

| Super family | (GPCRs) Biogenic amine receptors |  |  |
| --- | --- | --- | --- |
| Group | Member |  |  |
|  | NRNC Symbol | Name | Gene |
| adrenoreceptor | ADRA1A | adrenoceptor alpha 1A | ADRA1A |
|  | ADRA1B | adrenoceptor alpha 1B | ADRA1B |
|  | ADRA1D | Adrenoceptor alpha 1D | ADRA1D |
|  | ADRA2A | adrenoceptor alpha 2A | ADRA2A |
|  | ADRA2B | adrenoceptor alpha 2B | ADRA2B |
|  | ADRA2C | adrenoceptor alpha 2C | ADRA2C |
|  | ADRB1 | adrenoceptor beta 1 | ADRB1 |
|  | ADRB2 | adrenoceptor beta 2 | ADRB2 |
|  | ADRB3 | adrenoceptor beta 3 | ADRB3 |
| 5-hydroxytryptamine (serotonin)receptor | HTR1A | 5-hydroxytryptamine receptor 1A | HTR1A |
|  | HTR1B | 5-hydroxytryptamine receptor 1B | HTR1B |
|  | HTR1D | 5-hydroxytryptamine receptor 1D | HTR1D |
|  | HTR1E | 5-hydroxytryptamine receptor 1E | HTR1E |
|  | HTR1F | 5-hydroxytryptamine receptor 1F | HTR1F |
|  | HTR2A | 5-hydroxytryptamine receptor 2A | HTR2A |
|  | HTR2B | 5-hydroxytryptamine receptor 2B | HTR2B |
|  | HTR2C | 5-hydroxytryptamine receptor 2C | HTR2C |
|  | HTR4 | 5-hydroxytryptamine receptor 4 | HTR4 |
|  | HTR5 | 5-hydroxytryptamine receptor 5 | HTR5 |
|  | HTR6 | 5-hydroxytryptamine receptor 6 | HTR6 |
|  | HTR7 | 5-hydroxytryptamine receptor 7 | HTR7 |
| Dopamine receptor | DRD1 | dopamine receptor D1 | DRD1 |
|  | DRD2 | dopamine receptor D2 | DRD2 |
|  | DRD3 | dopamine receptor D3 | DRD3 |
|  | DRD4 | dopamine receptor D4 | DRD4 |
| Histamine receptor | HRH1 | Histamine receptor 1 | HRH1 |
|  | HRH2 | Histamine receptor 2 | HRH2 |
|  | HRH3 | Histamine receptor 3 | HRH3 |
|  | HRH4 | Histamine receptor 4 | HRH4 |
| Trace amine associated receptor | TAAR1 | trace amine associated receptor 1 | TAAR1 |
|  | TAAR2 | trace amine associated receptor2 | TAAR2 |
|  | TAAR3 | Trace amine associated receptor 3 | TAAR3 |
|  | TAAR4 | Trace amine associated receptor 4 | TAAR4 |
|  | TAAR5 | Trace amine associated receptor 5 | TAAR5 |
|  | TAAR6 | Trace amine associated receptor 6 | TAAR6 |
| Muscarinic Acetylcholine receptor | mAChR-A | muscarinic acetylcholine receptor A-type | mAChR-A |
|  | MAChR-B | muscarinic acetylcholine receptor B-type | MAChR-B |
|  | MAChR-C | muscarinic acetylcholine receptor C-type | MAChR-C |

The multiple sequence analysis was performed in all available genomes of the biogenic amine receptors family with putative full-length protein sequences. Based on results, the biogenic receptors members were identified in mammals and most of them in birds, fishes, lizards, amphibians and reptiles. Only few of the members were identified in plants (HTR2C, HTR4, HRH1, and HRH2) and in insects (ADRA2C, HTR1b, HTR2a, HTR2b, HRH1, H RH2, HRH3, mAChR, TAAR1, and TAAR6). We have to mention that adrenoceptors alpha- 1were identified only in *Homo sapiens* and *Mus musculus* species, since their sequences were recently uploaded. By contrast, 5- ht1A receptors were identified in mammals, birds, reptiles, fishes, amphibians and turtles (Table 2).

**Table 2.**
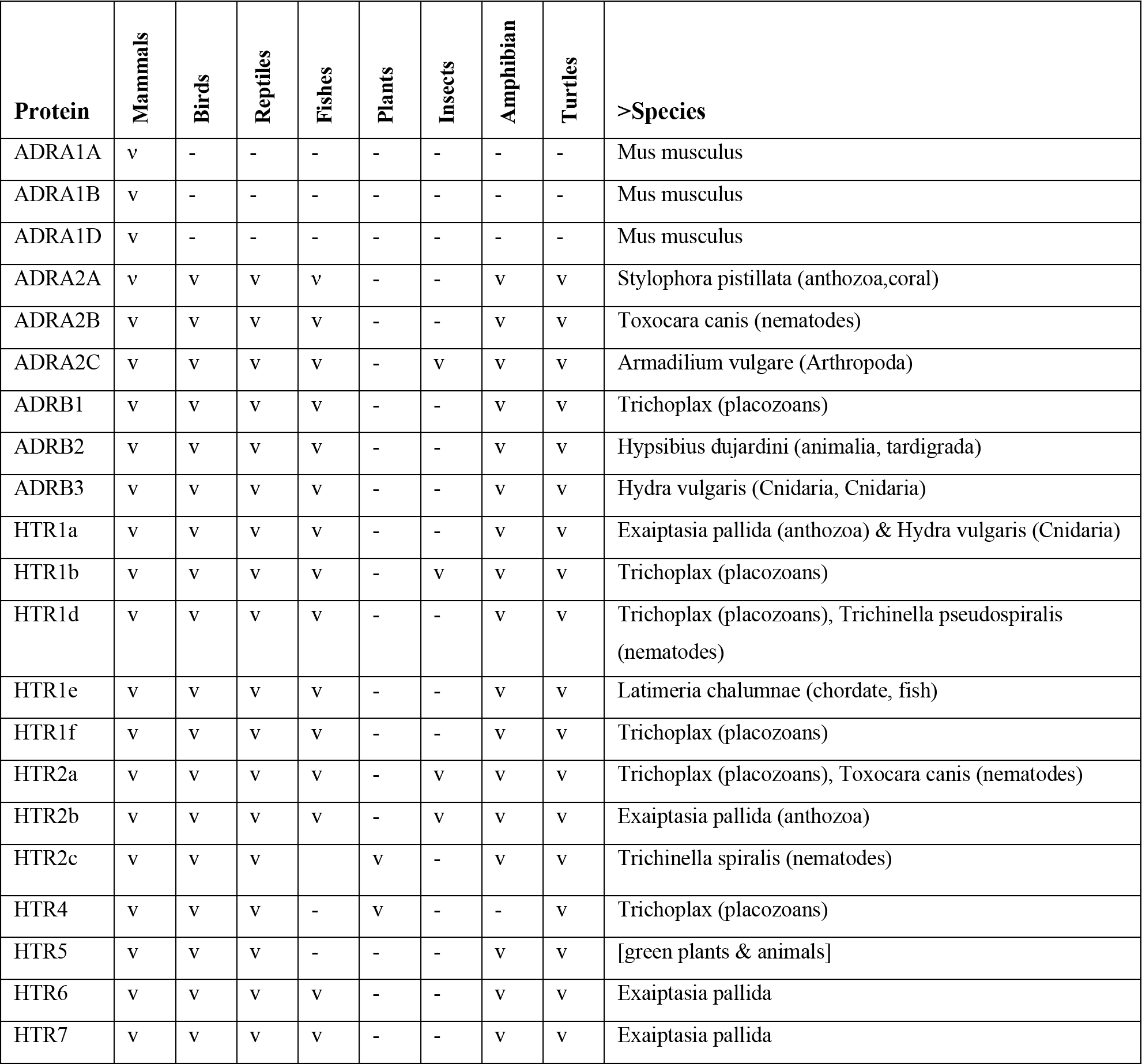
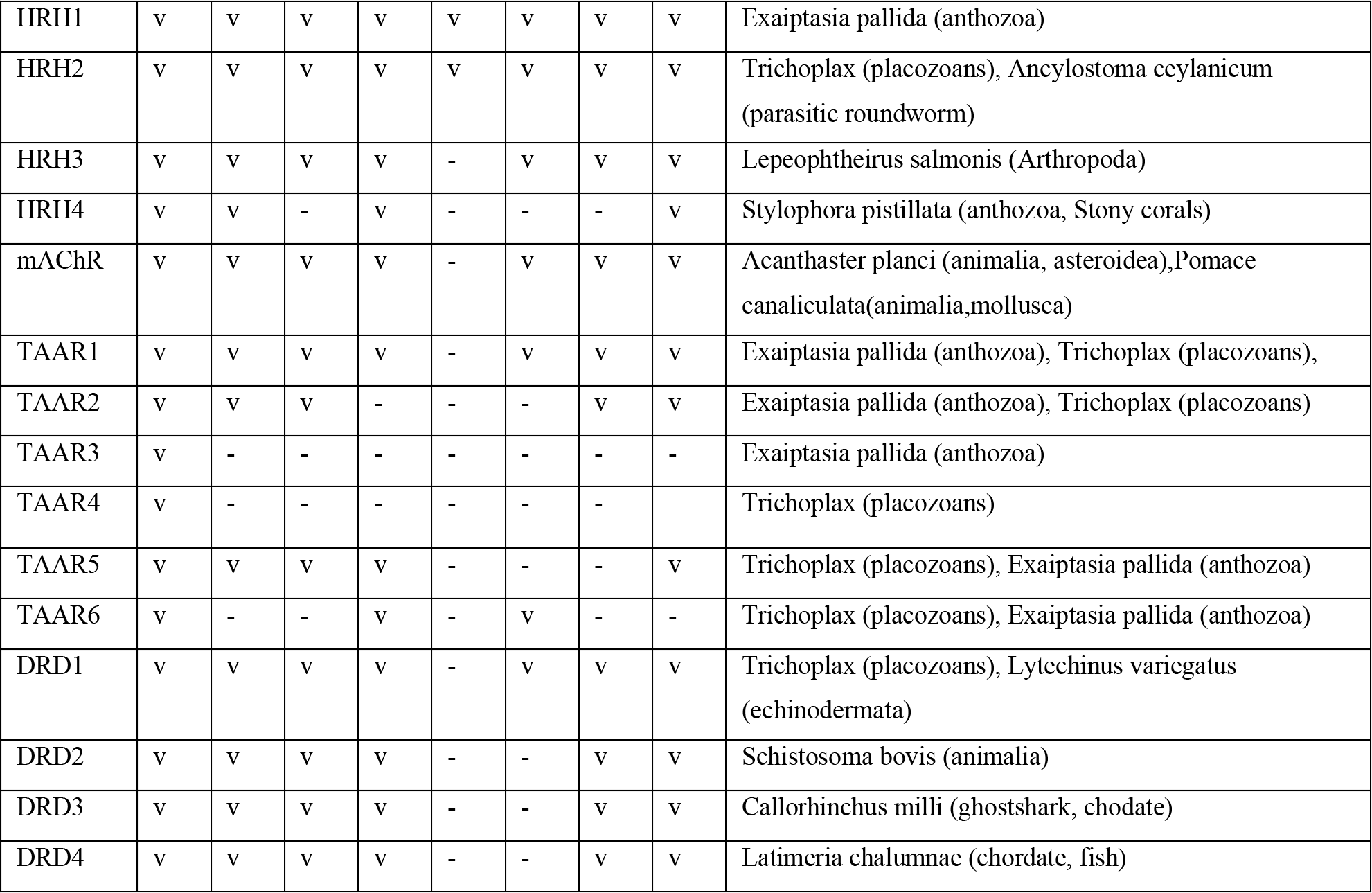
Categorization of the biogenic amine receptors members per cluster and phylum. In the last column are the representative organisms that correspond to the more distant species.

The GPCRs and biogenic amine receptors have a characteristic existence of seven membrane-spanning a-helical segments which are separated by alternating intracellular and extracellular loop regions (Figure 1 and 2). As seen in the 3D structure of bioamine receptors, the intracellular domain is arranged after the fifth a-helix, followed by the remaining two extracellular a-helices VI and VII. The main differences between these members, apart from the distribution of amino acids, are the intracellular domain and the loops that contain the transmembrane helices, especially at the edges and after the fifth a-helix (Figure 1 and 2).

### The conserved signaling motifs of the GPCRs

Multiple sequence alignment of the adrenoceptors, the 5-hydroxytryptamine receptors and in general, the biogenic amine receptor family’s protein sequences from a variety of several species were included in the first sub-tree, highlights important conserved functional domains. There is evidence indicating highly conserved domains throughout the length of the sequence which belongs to the transmembrane domain (helices 1-7), especially among species that belong to the same taxonomic division (Figure 4). An effort has been done in this study towards characterizing conserved motifs that play a key role in the biogenic amine receptor family.

**Figure 4.**
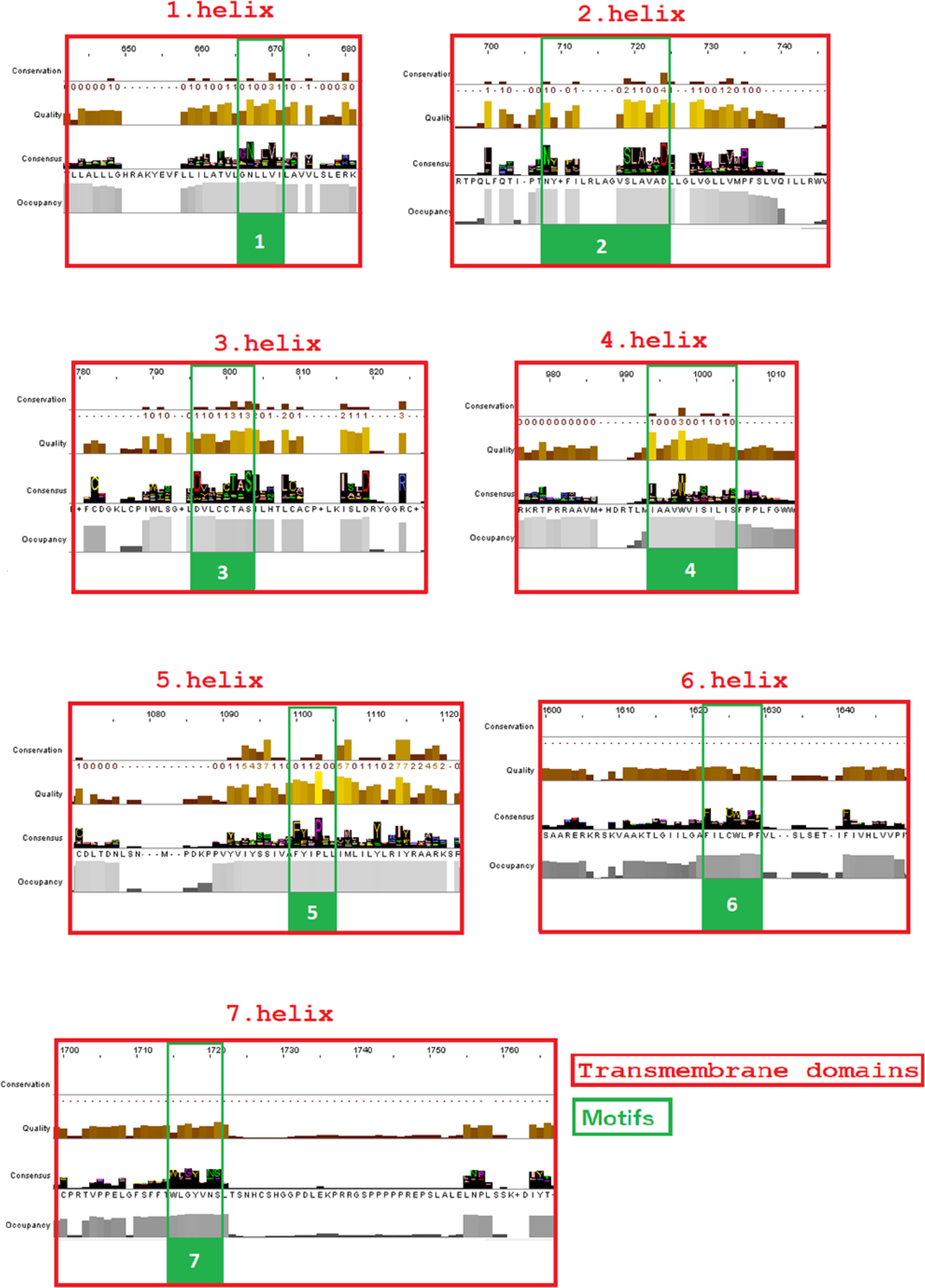
Highly conserved motifs in the biogenic amine receptors. Motifs 1 to 7 have been highlighted (colored green) in the intracellular domain, based on the consensus sequence from the MSA. All the motifs have been found in the transmembrane helices (colored red). The conserved motifs were identified through the consensus sequence and logo graph where generated using Jalview software.

Finally, the consensus sequence of the multiple sequence alignment highlights eleven conserved motifs between adrenoceptors, ten between 5-hydroxytryptamine receptors and seven which are conserved between all species of the biogenic amine receptor family (Figure 4). All of the conserved motifs identified have not been reported previously, apart from those which are located in helix 6 and helix 7 (intracellular domain), and indisputably deserve further study. It should be mentioned that the conserved motifs among species that belong to adrenoceptors are mainly at the edges of each intracellular helix, but among all the GPCR members of the biogenic amine family the key patterns in each intracellular helix have been identified in smaller patterns. The highly conserved motifs identified in the protein family are directly related to their active site (which is claimed to be a groove between helices III, IV, and V) and their protein function, since specific chemical compounds activate those GPCRs receptors towards transmitting biological messages between cells.

Regarding serotonin receptors, ten conserved patterns are identified. Note that HTR3 receptors were not included in the alignment as they do not belong to the GPCRs. It should be noted that the conserved motifs between members belonging to the serotonin receptors are located mainly within each a-helix, and extend between the transmembrane and intracellular domains. In this way they participate. In this way they are involved in both the formation of the ligand binding site and the binding of G proteins. All conserved patterns identified here have been previously mentioned, with the exception of helix 7 in which a more extensive pattern than the NPXXY of the literature was observed.

### Phylogenetic analysis

In the present study, we performed three representative phylogenetic analyses, related to the biogenic amine receptors family (GPCRs), the adrenoceptors sub-family and the 5-ht receptors sub-family, respectively. The phylogenetic reconstructions of the biogenic amine receptors and the adrenoceptors protein sequences indicate unambiguous clustering between the GPCR members as shown in Table 1 (Figure 3). Phylogenetic analyses topology and sub-clusters have been found to share similar organization based on the result from other studies (21, 56). Phylogenetic analyses executed by Henry Lin *et al*. and by Fredriksson *et al*. using a ligand- and sequence-based dendrograms show that we are corresponding at a high level (21, 56). In particular, our sub-clusters and GPCRs closely related the adrenoceptors and 5-ht receptors are conformed to these analyses. The biogenic amine receptors family’s phylogenetic tree contributed towards identifying which members are directly related with both ADRA2B and HTR1A during evolutionary progression. This correlation will help us to combine knowledge from other closely related GPCRs in order to study in more detail the mental disorders and in general the eating disorders as well as suicidal behavior. Based on results, adrenoceptors are closely related to the DRD2, DRD3, DRD4, TAAR, HTR5 and HTR1 receptors. Subsequently, 5-ht1 (HTR1) receptors are closely related to HTR5, DRD4, DRD3, DRD2 and ADRA receptors.

### Molecular modeling of the HTR1A intracellular domain

The structural comparisons between different 3D structures have been shown significant differences in the intracellular domain of each ADRA2B receptor and other members of biogenic amine receptors. In this study, there have been done many estimates of both secondary and third structure of this certain protein domain. Unfortunately, the results we got were not adequate, because information about the intracellular domain of the ADRA2B receptor is lacking. Up to now the more decent and detailed protein crystal of ADRA2B protein is the one with the accession number 6K41R which is from the *Bos taurus* organism, member of mammals.

Homologous solved 3D structures from the Protein Data Bank (PDB) have been identified from the Protein Data Bank (PDB) using the NCBI/BLASTp algorithm. The final choice of the template structure referring to HTR1A was not only based on the Percent Identity of the sequence and the analysis of the structure, but also on the results of the phylogenetic trees that were designed. Thus, in order to design the appropriate model for the 5 HTR1A, the following model templates were selected, PDB: 4IAR and 7E2X that meet specific criteria: Query Cover ≥ 90%, Percent Identity 30-80%, and after analysis the best cases were the complete crystal structure of the human receptor 5-HT1B (PDB: 4IAR) and the fragment structure of the 5-HT1A extracellular and transmembrane domains (PDB: 7E2X).

## Discussion

### Structural and functional description of ADRA2B and HTR1A

As predicted, from the sequence alignment of ADRA2B and its secondary structure data we investigated, the transmembrane and extracellular domains follow the structural features of known biogenic amine receptors (Figure 1 and 2). From the knowledge we have till now, the structure of ADRA2B receptor there are located four distinct domains, two of them are extracellular and the other two intracellular. The extracellular domains are the transmembrane helices and the extracellular loops (37), which are in a large extent maintained among biogenic amine receptors. On the other hand, the intracellular loops (ICL) and the intracellular domains are located into the cell and have many differences compared to the other members of the family. The intracellular domains of ADRA2B receptor are not analyzed enough in order to have a starkly secondary structure prediction. We guess that they have a combination of a-helices, beta sheets and some parts have not a certain conformation.

The transmembrane a-helices are seven in number and are a characteristic domain of all GPCRs (Figure 5 and 6). They are separated by alternating intracellular and extracellular loop regions with the first one being an ICL. As described by Cherezov 2007 helices II, V, VI, and VII each have a proline-induced twist at conserved positions along the span of the transmembrane segments (22). These twists are thought to enable the structural rearrangements required for activation of G protein effectors. The extracellular loops and N-termini of GPCRs, together with the extracellular halves of the transmembrane helices, are believed to define the ligand-binding site of each receptor. During activation, like all GPCRs, ADRA2B and HTR1A open a cavity on its cytoplasmic side, between the fifth and the sixth helix and the remaining of the receptor, to interact with downstream effectors.

**Figure 5.**
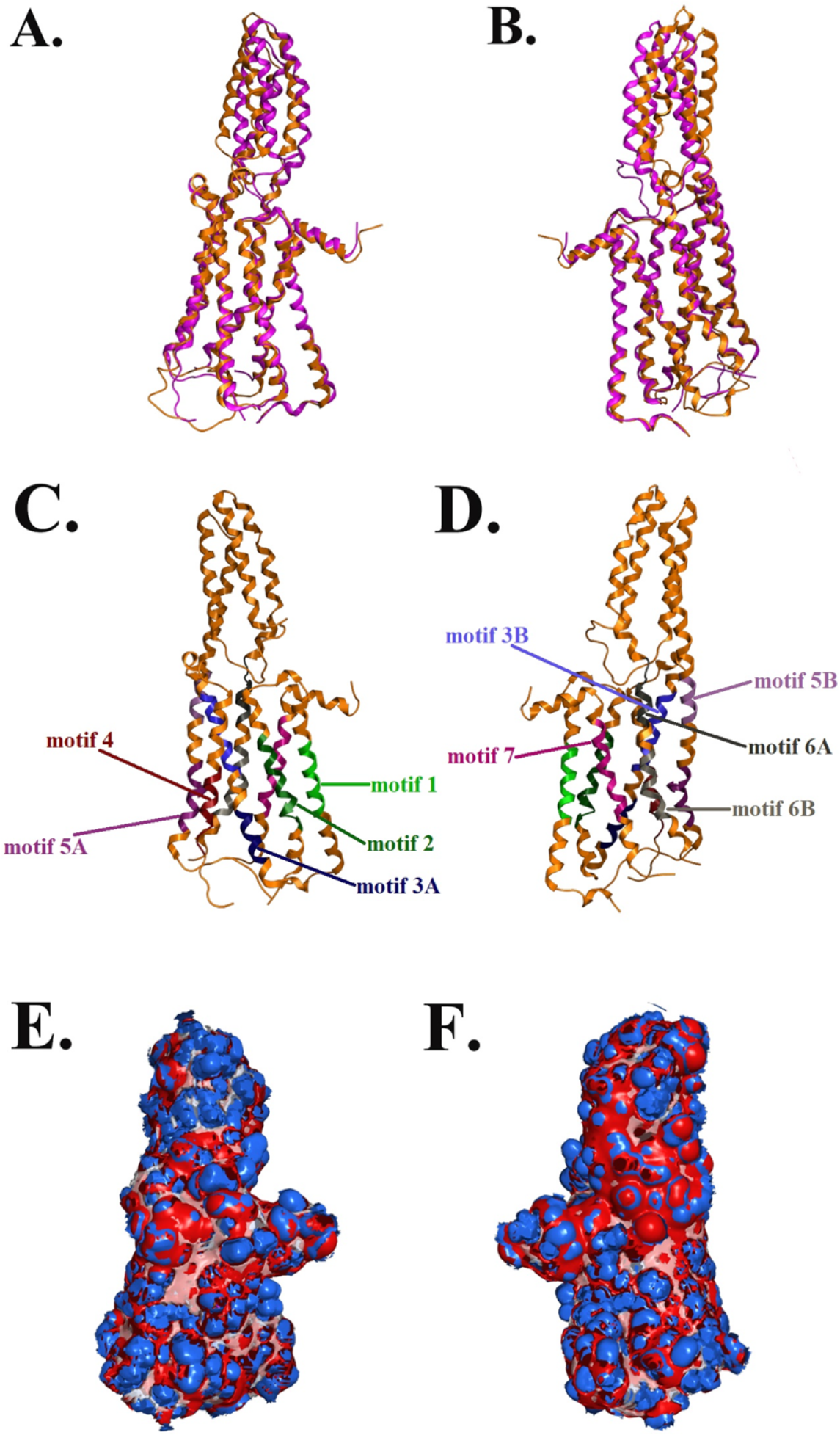
Homology model of the human 5-HT1A. (A-Top and B-Front) Ribbon representation of the produced human 5-HTR1A model (colored purple) superposed with the corresponding human HTR1B (in orange). (C - Top and D - Front) The ten suggested conserved motifs of the human 5-HTR1A. Electrostatic surface potential for the human 5-HTR1A. Represented with blue is the area of negative charge. Red is the area of positive charge and white is the un-charged region.

**Figure 6.**
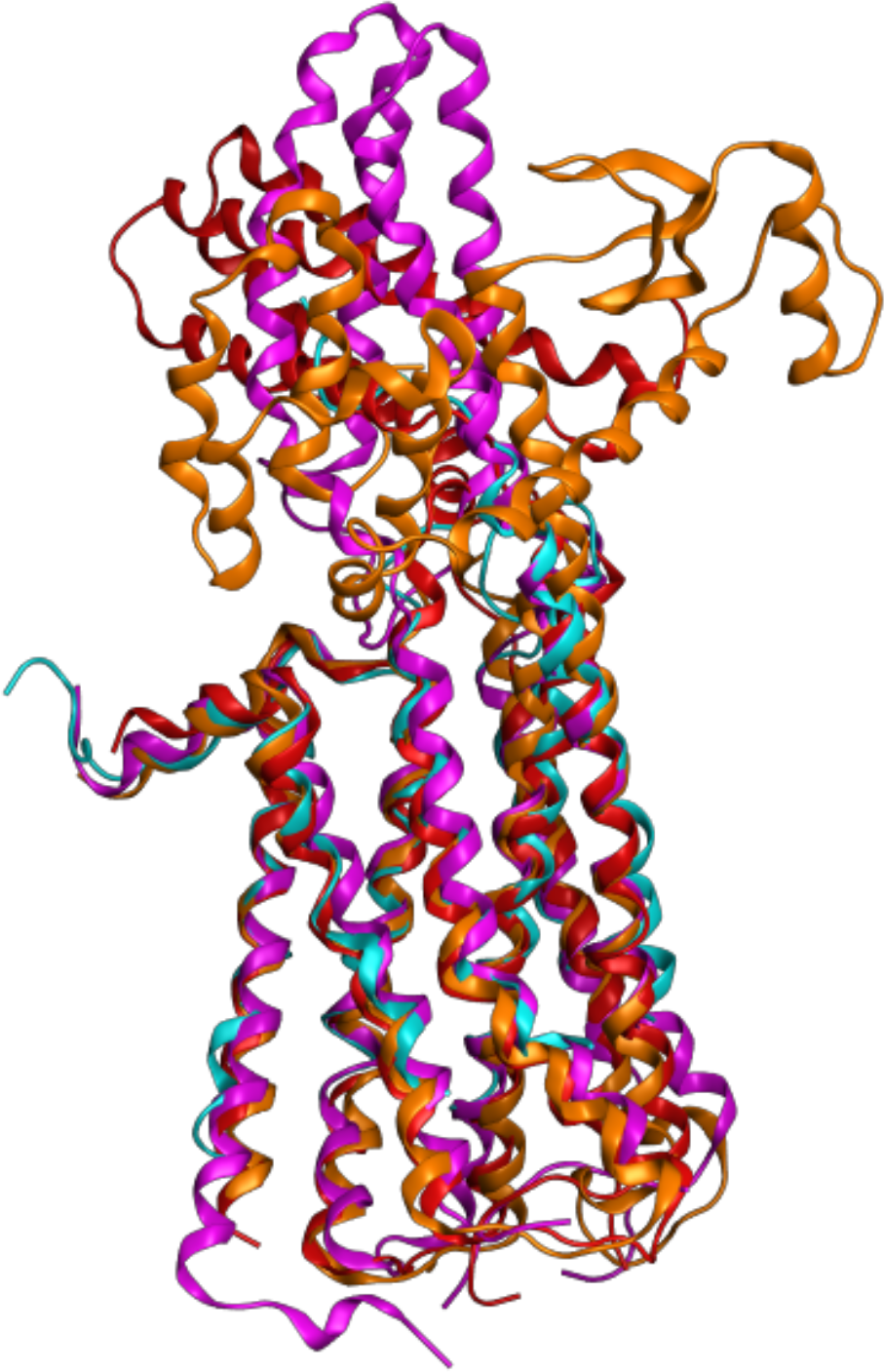
Structural superpositions of representative biogenic amine receptors in different poses. In the ribbon representation are shown the HTR1B (colored blue, 4IAR_A), the 5-hydroxytryptamine receptor 1A (5-HTR1A) (colored red | 7E2X_R), the DRD2 (colored purple | 6CM4_A) and the ADRA2A (colored orange | 6KUY_A).

Several studies have been shown that the dexmedetomidine binding pocket has similar positions in the formed orthosteric binding pockets for their agonist, but they differ in the actual types of interactions (ADRA2B is bound to dexmedetomidine and ADRB2 is bound to adrenaline) (57). In the case of β_2_AR, adrenaline binds primarily through an ionic interaction and hydrogen bonds with D113, S203, S207, N293 and N312. On the other hand, dexmedetomidine occupies a slightly deeper position in the binding pocket and interacts with the α_2_BAR primarily through aromatic and van der Waals interactions, besides the conserved ionic interaction to D92. As regards the structural basis of ADRA2B selectivity, it is claimed that of the 13 amino acids that construct the binding pocket of adrenaline in ADRB2, 4 of them are different in ADRA2B, which are C96, L166^4^, Y391 and F412. From previous studies it is revealed that F412 in ADRA2B is essential for binding a2-selective imidazoles like dexmedetomidine. The resolved structure of α2BAR revealed that F412 is responsible for the formation of an enclosed sub-pocket that binds the positively charged imidazole ring of dexmedetomidine (57). When adrenaline is attached to ADRA2B addresses the same pocket with its methylamino group. Although, this does not happen in case for isoprenaline group but it leads to repulsive interactions. That is why the selectivity of ADRB agonist is oprenaline over ADRA2B. From sequence alignment of all adrenoceptors it is revealed that the amino acids which take place to the dexmedetomidine binding pocket in ADRA2B are conserved in all α2-adrenergic receptors subtypes. The only difference is that ADRA2A has I45 in ECL2 but both ADRA2B and ADRA2C have L45. A definite characteristic of all subtypes of ADRA2 pockets is Y391 that can’t be found in any other adrenergic receptor. In the active structure of ADRA2B, Y391 interacts with the phenyl ring of dexmedetomidine and forms a hydrogen bond with S176 in TM5.

Even though there is no predicted inactive state of ADRA2B available, Daopeng Yuan *et al*. performed molecular dynamics simulations of ADRA2B after removing Go and dexmedetomidine from the complex (57). As they found out, TM5 is displaced inward and there is a right-handed rotation of TM6 and TM7. So, maybe interactions between the four aromatic residues on TM6 and the phenyl and imidazole rings of dexmedetomidine lead to a rotation of TM6, rearrangement of the PI F motif and outward displacement of the cytoplasmic end of TM6.The interactions between ADRA2B and Go proteins are almost 17 in number and located mainly at ICL1, ICL2, TM5, TM4, TM3 and TM6 (52).

The conserved motifs located in the transmembrane helices play an important role in the agonist-binding affinity and the activation of the receptor (Figure 5 and 6). It should be mentioned that among biogenic amine receptors we detected the existence of cysteines that are preserved among all members of the family. These cysteines are located at the extracellular loops before the third and the fifth helices accordingly (58). In general, these cysteines exist because they can be connected together by a disulfide bond that helps to stabilize the structure of a molecule; in this case, they stabilize the exposed extracellular domains.

Another notable comment is that there is a poly-E domain in the sequence of ADRA2B that belongs to the intracellular domain of the protein (298–309), especially at the third intracellular loop. Since a deletion variant of three of these glutamic acid residues (301–303) increases noradrenaline availability, this poly-glutamic acid domain may be important for the noradrenaline expression and the function of ADRA2B (38, 39). Glutamic acid is an acidic amino acid found in proteins that have a prominent role in enzyme active centers and in maintaining the solubility and ionic character of proteins (59). Glutamic acids are mainly located to the protein surfaces so they have access to the solvent and is more likely to be found in helices (59). The carboxylate anions and salts of glutamic acid are known as glutamates. Glutamate is known to follow up on a few distinct sorts of receptors and has excitatory impacts at ionotropic receptors and modulatory impacts at metabotropic receptors [which are G protein-coupled glutamate receptors (mGluR) that adjust neuronal and glial sensitivity through G protein subunits acting on membrane ion channels and second messengers such as diacylglycerol and cAMP (59). Glutamate plays an important role in synaptic plasticity in the brain and is involved in various cognitive functions, such as learning and memory. As Haber-Pohlmeier *et al*. describe at their article, glutamic acid-rich proteins in vertebrate rod photoreceptors may be loose coils with low-affinity but high-capacity Ca2+ binding (60).

### Description of the 5-hydroxytryptamine receptor 1A intracellular domain model

From the information obtained about the crystal structure of the human 5 HT1B receptor in complex with ergotamine, a migraine-fighting drug, a large ligand-binding cavity was revealed, defined by residues of helices III, V, VI, VII and ECL2, comprising an orthosteric pocket embedded deep in the 7TM core and an extended binding pocket near the extracellular entrance (61). The same formation also found in the crystal structure of the 5-hydroxytryptamine receptor 1A (5-HTR1A) (PDB: 7E2X_R) extracellular and transmembrane domains (53) (Figure 6).

Helix movements during receptor activation are accompanied by a common set of local “microswitches” in the intracellular domain of GPCRs. The microswitches are characterized by configuration changes in highly conserved side chains, creating rotamers, which stabilize the helix movements and help prepare the intracellular side of the GPCR for G protein binding(62). GPCR activation is an allosteric process that links agonist binding and G protein uptake to the characteristic external movement of transmembrane helix 6 (TM6). This movement is a common feature of receptor activation; however, changes in the level of residues that cause the movement of TM6 remain less understood. One of the most prominent examples of ligand - dependent rearrangement at the binding pocket involves a shift of the Trp6.48 residue. Although this side chain was previously classified as a microswitch (or “rotamer toggle switch”), the crystal structures of the active GPCR states revealed no changes in this residue upon activation. Nevertheless, the crystal structures suggest a prominent role for Trp6.48 as a significant ligand-dependent ‘trigger’ in some GPCRs (62, 63). The conserved motifs are located in the helices of the transmembrane regions and play an important role in the activation of the receptor and in the binding of the agonist molecules that interact with these proteins (Figure 5 and 6).

The D[E]RY motif at the cytoplasmic terminus of helix III and intracellular loop 2 (ICL2) represents one of the most conserved motifs of GPCR class A, rich in Arg^3.50^ residues (96% conservation among GPCR A) (64). It forms a salt bridge with the adjacent acidic side chain Asp (Glu) 3.49 (Asp 68%, Glu 20%) and with TM6, known as the ionic lock that limits the receptor to an inactive configuration. Once the ligand binds, the ionic lock breaks and DRY forms new interactions with TM5, stabilizing the receptor in an active configuration (62, 65). The NPxxY motif is located near the intracellular end of helix VII and contains a highly (92%) conserved Tyr7.53 residue that serves as an important activation microswitch in GPCRs. In inactive GPCR structures, the side chain of Tyr7.53 turns towards helices I, II or VIII. In contrast, in all GPCR active state crystal structures, the Tyr7.53 side chain changes its configuration (rotamer) and rotates toward the middle axis of the 7TM beam, forming interactions with the side chains of helices VI and III (62). The CWxP pattern in TM6 (6.47 6.50) is largely maintained in non-olfactory GPCR class A, with C6.47, W6.48 and P6.50 present in 71%, 78% and 98% of sequences, respectively. P6.50 creates a huge twist on the TM6 helix (causing a helix rotation of about 35°), much larger than an ordinary proline in a transmembrane helix (about 20°).

Various amino acid substitutions in the transmembrane regions Asp-82 to Asn-82, Asp-116 to Asn-116 and Ser-198 to Ala-198 have resulted in a decrease in affinity for 5-HT by 100 fold. Mutant Thr-199 to Ala-199 showed virtually no binding. Further mutation studies revealed that Ser-392 and Asn-395 on may also be crucial for ligand binding (66). Amino acid Asp-82 is found in TM-2 and specifically within the motif 2A, Asp-116is found in TM-3 and specifically within the motif 3A. Both Ala-198 and Thr-199 are placed in motif 5A of TM-5. Amino acids Ser-392 and Asn-395 of TM-7 are located in motif 7.

The initial breakthrough concerning the 5-HTR1A ligands were the identification of 8-OH-DPAT, a selective 5-HT1A receptor agonist, during the 80s. It’s application provided the first pharmacological profile of the 5-HT1A binding site. It was also shown that buspirone and a series of structurally related 5-HT1A ligands were anxiolytic and antidepressant in the clinic. Following these discoveries, several tryptamine derivatives have followed. Carboxamidotryptamine (5-CT) have demonstrated affinity for 5-HT1A receptors similar to that of serotonin. It also seemed that the conformational restricted tryptamine analogue RU28253 has a good affinity for the 5-HT1A receptor and indicated greater affinity for the 5-HT1B sites. Gepirone is a structural analogue of the non-benzodiazepine anxiolytic buspirone and belongs to the long-chain arylpiperazine (LCAP) 5-HT1A receptor ligands. Gepirone possesses a much greater selectivity for the 5-HT1A receptor over the 5-HT2A and dopamine D2 sites. At this point, the issue of non-selective binding arises. Well utilized 5-HT1A receptor agonists, such as buspirone, tandospirone and ipsapirone show poor selectivity for 5-HT1A receptor. Buspirone and tandospirone exhibit affinity for the dopamine D2 receptor and ipsapirone shows affinity for the α1-adrenergic receptor. Non-selective 5-HT1A receptor drugs can act as dopamine D2 antagonists and may cause side effects such as prolactin stimulation. The selectivity of the 5-HT1A receptor over the dopamine D2 and α1-adrenergic receptors seem to be due to structural differences and it is highly probable that these differences are the key to developing alternative therapies. The 5-HT1A receptor agonists are valuable tools to develop treatments for anxiety and depression (66, 67).

## Conclusion

Despite the enormous scientific interest in GPCRs, and biogenic amine receptors family, current knowledge of their evolutionary analyses and their function is still quite limited. The highly conserved motifs for these proteins could be used as promising targets for a more specialized characterization of this huge protein family. These motifs could be used as innovative identification targets, through the development of new technologies concerning the numerous critical pathways, affected by this protein family. Considering the phylogenetic trees in the present study, useful beneficial insights are provided for the biogenic amine receptors family’s evolution. As concerns the ADRA2B receptor we gathered much information in order to characterize its structure and function, even though the data for this receptor is limited. The comparison between the HTR1A and ADRA2B 3D models the pathways of these receptors and in an effort to discover more clues about the structure-function relationship.

## Abbreviations

GPCR: G protein-coupled receptor
MOE: Molecular Operating Environment
ECL: Extracellular loops
ICL: Intracellular loops
TM: Transmembranic helix
EBP: Extended Binding Pocket EI Emotional Intelligence
SNPs: Single Nucleotide Polymorphisms Matlab Matrix Laboratory
MEGA: Molecular Evolutionary Genetics Analysis
NCBI: National Center for Biotechnology Information

## Acknowledgments

Not applicable.

